# Identification and experimental verification of key genes related to the Ras signaling pathway and the Hippo signaling pathway in osteoarthritis based on transcriptome data

**DOI:** 10.64898/2026.02.09.704991

**Authors:** Lina Zhang, Ying Lu, Diefeng Liu, Bin Sheng

## Abstract

**Background:** Osteoarthritis (OA) is a chronic degenerative joint disease characterized by the progressive deterioration of articular cartilage, significantly impacting the quality of life in middle-aged and elderly populations. The Ras and Hippo signaling pathways play critical roles in regulating cell proliferation, differentiation, and stress responses; however, their interactive mechanisms in OA remain unclear. This study aimed to identify key genes associated with these two pathways using bioinformatic approaches and to elucidate their potential mechanisms in OA.

**Methods:** Transcriptomic data of OA along with Ras signaling pathway-related genes (RSPRGs) and Hippo signaling pathway-related genes (HSPRGs) were obtained from public databases. Differentially expressed genes (DEGs) were identified, and key genes were screened through machine learning, expression validation, and receiver operating characteristic (ROC) curve analysis. Functional insights were further explored via gene set enrichment analysis (GSEA), subcellular localization, immune infiltration analysis, regulatory network construction, and drug prediction. Finally, the expression of key genes was validated in clinical samples.

**Results:** KIT and CSF1R were identified as key genes. GSEA indicated their involvement in pathways such as the lysosome pathway. Subcellular localization predicted that KIT and CSF1R are distributed in the nucleus, extracellular region, and plasma membrane. Immune infiltration analysis revealed that KIT showed a positive correlation with eosinophils and a negative correlation with immature dendritic cells, while CSF1R was positively correlated with macrophages and negatively correlated with CD56ᵇʳⁱᵍʰᵗ natural killer cells. Drug prediction suggested interactions between the key genes and several therapeutic agents, including avapritinib and IMC-CS4. Subsequently, we validated our findings in cartilage tissue samples and discovered that compared to the control group, both CSF1R mRNA and protein expression was significantly upregulated in OA tissue, while KIT expression was significantly downregulated.The same results were also validated in immunofluorescence staining of chondrocytes.

**Conclusion:** This study identified KIT and CSF1R as key genes in OA, providing new theoretical insights and potential targets for mechanistic research and targeted therapy.

## Introduction

Osteoarthritis (OA) is a chronic, degenerative joint disorder characterized by progressive articular cartilage loss, subchondral bone sclerosis, osteophyte formation, synovial inflammation, and periarticular muscle atrophy[1]. Globally, OA represents the leading cause of pain and disability among elderly populations, with its prevalence increasing markedly with age. In the context of rapid global population aging, OA has become a major public health concern, imposing significant socioeconomic and healthcare burdens worldwide [2]. The clinical diagnosis of knee OA currently relies primarily on patient symptoms and imaging evaluations[3]. However, conventional imaging modalities such as radiography can only detect advanced structural alterations, demonstrating limited sensitivity for early-stage disease detection. As a result, many patients are diagnosed only after irreversible cartilage damage has occurred. Current therapeutic strategies largely focus on alleviating pain and inflammation but remain insufficient to prevent or reverse cartilage degeneration during the early course of OA [4]. Therefore, elucidating the molecular mechanisms underlying OA pathogenesis and identifying pivotal regulatory genes are critical for the development of early diagnostic biomarkers and effective disease-modifying treatments.

The pathogenesis of osteoarthritis involves dysregulation across multiple signaling pathways, with the inflammatory cascade and abnormal mechanical stress perception playing central roles in cartilage degeneration. The RAS signaling pathway, named after its central GTPase RAS, is a canonical intracellular cascade that transduces extracellular growth factor signals through the sequential activation of RAF, MEK, and ERK kinases. This pathway regulates essential cellular processes including proliferation, differentiation, apoptosis, and inflammatory responses[5,6]. Recent studies have demonstrated that during OA progression, activation of the RAS-ERK axis promotes synovial inflammation and enhances the production of proinflammatory mediators such as interleukin-1β (IL-1β) and tumor necrosis factor-α (TNF-α). TNF-α, in turn, induces chondrocyte apoptosis through dual activation of chondrocytes and synovial fibroblasts, thereby amplifying inflammation and accelerating matrix degradation. This establishes a self-sustaining inflammatory feedback loop characterized by continuous release of catabolic enzymes and cytokines, ultimately exacerbating cartilage breakdown and pain sensitization.

The Hippo signaling pathway, an evolutionarily conserved regulator of tissue growth and homeostasis, operates through the MST1/2–LATS1/2 kinase cascade, which restricts the nuclear translocation of the transcriptional coactivators YAP and TAZ. Through this mechanism, Hippo signaling suppresses proliferative gene expression and maintains cellular equilibrium [7,8]. In articular cartilage, the Hippo-YAP/TAZ axis functions as a key mechanosensitive regulator, integrating extracellular mechanical cues to control chondrocyte proliferation, differentiation, and matrix synthesis. Physiological mechanical loading inhibits Hippo signaling to activate YAP/TAZ, promoting cartilage repair and regeneration, whereas abnormal mechanical stress disturbs this balance, leading to chondrocyte dysfunction and OA progression. Moreover, YAP/TAZ activity has been closely associated with the regenerative potential of cartilage-resident stem and progenitor cells [9]. Notably, functional crosstalk exists between the RAS and Hippo pathways. Oncogenic RAS can suppress Hippo signaling to activate YAP/TAZ, suggesting that these two pathways may cooperatively regulate chondrocyte fate and joint homeostasis [10].

Current research primarily focuses on the roles of individual genes within single pathways in OA, lacking systematic investigations into the overall expression patterns, interactive regulatory networks, and diagnostic/therapeutic target potential of genes associated with the RAS and Hippo pathways in OA. The association between these pathway genes and the OA immune microenvironment also remains poorly elucidated. Therefore in this study, transcriptomic datasets related to OA and genes involved in RAS and Hippo signaling were obtained from public databases. Differentially expressed genes were identified and subsequently refined through machine learning algorithms, expression validation, and receiver operating characteristic (ROC) analysis to determine key OA-associated genes. Comprehensive bioinformatics analyses were then conducted to elucidate the molecular functions, signaling pathways, and immune microenvironment characteristics associated with these genes, providing novel insights into OA pathogenesis and potential targets for early diagnosis and precision therapy.

## 2. Materials and methods

### 2.1 Data collection

The datasets GSE114007 and GSE57218 were downloaded from the Gene Expression Omnibus (GEO, https://www.ncbi.nlm.nih.gov/geo/) database. The GSE114007 dataset, which was generated using the GPL11154 platform for high-throughput sequencing, served as the training set and included 10 cartilage tissue samples from patients with osteoarthritis (OA) and 10 normal cartilage tissue samples. This dataset was used for differential gene expression analysis. The GSE57218 dataset, based on the GPL6947 microarray platform, was employed as an independent validation set and consisted of 33 OA cartilage tissue samples and 7 normal cartilage tissue samples. It was utilized for external validation of expression results and receiver operating characteristic (ROC) analysis. Additionally, 238 Ras signaling pathway-related genes (RSPRGs) were retrieved from the Kyoto Encyclopedia of Genes and Genomes (KEGG, https://www.genome.jp/kegg/) using the keyword “Ras signaling pathway” (**Supplementary Table1)**. Meanwhile, 202 Hippo signaling pathway-related genes (HSPRGs) were obtained from the MsigDB database (https://www.gsea-msigdb.org/gsea/msigdb/) under the query term “Hippo” with the following filters: “H: hallmark gene sets”, “C2: curated gene sets”, “C5: ontology gene sets”, and “Homo sapiens” (**Supplementary Table2)**.

### 2.2 Identification of candidate genes

Differential expression analysis between OA and control samples (OA vs control) in the training dataset was performed using the “DESeq2” package (v 1.40.2) (PMID: 25516281) (|log_2_FoldChange (FC)| > 0.5 and P < 0.05). Volcano plots illustrating the differentially expressed genes (DEGs) were generated using the “ggplot2” package (v 3.5.1)[11], with the top 10 up-regulated and down-regulated genes labeled based on their log_2_FC values. Additionally, a heatmap was constructed using the “ComplexHeatmap” package (v 2.14.0)[12] to visualize the expression patterns of the top 10 up- and down-DEGs in OA and control groups. The “ggvenn” package [13] was employed to identify overlaps among the DEGs, RSPRGs, and HSPRGs, and a Venn diagram was plotted to screen candidate genes.

### 2.3 Functional analysis of candidate genes and protein-protein interaction (PPI) network construction

To investigate the biological pathways associated with candidate genes, Gene Ontology (GO) was performed on candidate genes using the “clusterProfiler” package (v 4.7.1.003) [14] and “GOplot” package (v 1.0.2) [15] (P.adj < 0.05). The top 3 enriched pathways from the GO categories of molecular function (MF), cellular component (CC), and biological process (BP) were selected and ranked based on their adjusted P.values (P.adj). KEGG pathway enrichment analysis was subsequently conducted using “clusterProfiler” package (v 4.7.1.003) for candidate genes, top 10 most significantly enriched pathways were visualized (P.adj < 0.05). To investigate the interactions among proteins encoded by candidate genes, PPI analysis was conducted using the Search Tool for the Retrieval of Interacting Genes/Proteins (STRING) platform (https://www.string-db.org/) with an interaction score threshold of > 0.4. The resulting PPI network was visualized using Cytoscape (v 3.10.2) [16].

### 2.4 Machine learning screening for candidate key genes

To further identify candidate key genes, least absolute shrinkage and selection operator (LASSO) regression analysis was applied to the candidate genes using the “glmnet” package (v 4.1-8) [17]. The optimal regularization parameter lambda was determined via 5-fold cross-validation using the cv.glmnet function. A model performance curve across lambda values was plotted, and the lambda value corresponding to the minimum point on the curve was selected to identify the LASSO-feature genes. The support vector machine-recursive feature elimination (SVM-RFE) algorithm was implemented with the “caret” package (v 6.0-94) [18], using 5-fold cross-validation to evaluate prediction accuracy across iterative feature subsets. The subset achieving the highest accuracy at the earliest iteration was selected, and the corresponding genes were retained as SVM-RFE-feature genes. For the eXtreme Gradient Boosting (XGBoost) algorithm, a binary classification model with a binary logistic objective function was constructed using the “xgboost” package (v 1.7.10.1) [19]. Hyperparameter tuning was performed via grid search and cross-validation. After model training, the importance of each candidate gene was assessed using the xgb. An importance function based on the Gain metric, and a feature importance bar plot were generated to select the XGBoost-feature genes. Finally, the “ggvenn” package (v 1.7.3) was used to identify candidate key genes among the feature genes obtained from 3 machine learning algorithms.

### 2.5 The correlation between Yes-associated protein (YAP) and candidate key genes

To investigate the potential interactions between YAP and the proteins encoded by the candidate key genes, PPI analysis was performed using the STRING platform (https://www.string-db.org/). The analysis included the candidate key genes and YAP1, with an interaction score threshold set to greater than 0.4. The resulting PPI network was visualized using Cytoscape (v 3.10.2). Subsequently, correlation analysis was performed using the “psych” package (v 2.4.3) (PMID: 37505622) to evaluate the relationships between the candidate key genes and YAP1 (|correlation coefficients (cor)| > 0.3, p < 0.05).

### 2.6 Expression level validation and ROC analysis

To validate whether the expression trends of candidate key genes were consistent between the training set GSE114007 and validation set GSE57218, the Wilcoxon test was employed to compare their expression levels between OA and control groups (P < 0.05). The results were visualized using violin plots generated with the “ggplot2” package (v 3.5.1). To further evaluate the ability of candidate key genes to discriminate between OA and control groups, ROC analysis was performed on the candidate key genes using the “pROC” package (v 1.18.5) [20] in the training dataset. The area under the curve (AUC) values were calculated, and candidate key genes demonstrating AUC values > 0.7 were identified as key genes.

### 2.7 Construction and evaluation of the nomogram model

To evaluate the predictive value of the key genes for OA, a nomogram model was constructed based on the training dataset using the “rms” package (v 6.5.0) [21]. The calibration curve, also generated with the “rms” package (v 6.5.0), was applied to assess the predictive accuracy of the nomogram within the training set, and the model fit was further evaluated using the Hosmer-Lemeshow test (HL test). To validate the predictive performance of the nomogram, ROC analysis was performed and the AUC value was calculated for the training set using the “pROC” package (v 1.18.5) (AUC > 0.7). Finally, the clinical utility of the nomogram was illustrated using decision curve analysis (DCA) implemented via the “rmda” package (v 1.6) [22].

### 2.8 GeneMANIA, chromosome localization, and correlation analysis of key genes

To investigate the interactions and functional associations between the key genes and genes with similar roles, a gene network was constructed by inputting the key genes into the GeneMANIA database (http://www.genemania.org). The chromosomal locations of the key genes were identified, and their distribution across chromosomes was visualized using the “RCircos” package (v 1.2.2) [23]. To analyze correlations among these biomarkers, Spearman correlation analysis was performed on the expression levels of the key genes in the training dataset using the “psych” package (v 2.4.3).

### 2.9 Gene set enrichment analysis (GSEA) and subcellular localization

To evaluate the enrichment of key genes in signaling pathways, GSEA was performed using the “clusterProfiler” package (v 4.7.1.003) to identify regulatory pathways or biological functions associated with key gene expression. Gene sets from the C2 collection of MSigDB database (https://www.gsea-msigdb.org/gsea/msigdb/), specifically the KEGG pathway subset, were retrieved using the “msigdbr” package (v 7.5.1) [24] with the species set to “Homo sapiens“, category to “C2”, and subcategory to “CP: KEGG”. Spearman correlation coefficients between key genes and all other genes were calculated using the “psych” package (v 2.2.9) [25], followed by ranking from high to low (|normalized enrichment score (NES)| > 1, P < 0.05, and P. adj < 0.25). Additionally, subcellular localization information for the key genes was retrieved from the GeneCards database (https://www.genecards.org/) to illustrate their distribution across cellular compartments.

### 2.10 Immune infiltration

To characterize the infiltration abundance of immune cells between OA and control samples in the training set, enrichment scores for 28 immune cell types [26] were calculated using the single-sample gene set enrichment analysis (ssGSEA) algorithm implemented in “GSVA” package (v 1.46.0) [27]. The resulting immune cell infiltration profiles were visualized using “ggplot2” package (v 3.5.1). Wilcoxon rank-sum tests were subsequently applied to compare immune cell infiltration levels between the OA and control groups. The results were presented as boxplots generated with “ggplot2” package (v 3.5.1). Immune cell types showing significant differences in infiltration abundance were defined as differential immune cells (P < 0.05). In the training dataset, spearman correlation analysis was performed using the “psych” package (v 2.2.9) to investigate both the relationships between differential immune cells and the associations of key genes with differential immune cells (|cor| > 0.3, p < 0.05).

### 2.11 Construction of transcription factor (TF)-microRNA (miRNA)-mRNA regulatory network

To elucidate the molecular regulatory mechanisms of key genes, an interaction network was constructed using NetworkAnalyst (https://www.networkanalyst.ca/). On the platform, “Gene list input” was selected with the species set to Homo sapiens and the input ID type specified as Official Gene Symbol. The key genes were entered into the gene list field and uploaded. The “TF-miRNA regulatory Network” analysis option was then chosen, which utilizes data from the RegNetwork database (http://www.zpliulab.cn/RegNetwork/). After proceeding, the resulting TF-miRNA-mRNA regulatory network was downloaded and imported into Cytoscape (v 3.10.2) for visualization.

### 2.12 Drug prediction for OA

To predict potential therapeutic drugs for OA, the DGIdb database (https://www.dgidb.org/) was used to identify potential drugs or molecular compounds interacting with the key genes. Subsequently, a drug-key gene interaction network was constructed using Cytoscape (v 3.10.2).

### 2.13 Cartilage sample collection

OA cartilage samples (n=5) were obtained from patients who underwent total knee arthroplasty (TKA) with a clinical diagnosis meeting the criteria for knee osteoarthritis[28] and an X-ray image diagnosis meeting the Kellgren-Lawrence classification grade IV[29]. The OA cartilage samples used in this study were all obtained from injured cartilage tissue at the edge of the cartilage full-thickness defect in the weight-bearing area of the medial tibial plateau. Non-OA cartilage samples (n=5) were obtained from knees without OA in patients with traumatic lower extremity amputation. The study protocol received ethical approval from the Institutional Review Board of Hunan Provincial People’s Hospital (**Approval No.[2025]-330**).

### 2.14 Reverse transcription quantitative PCR (RT-qPCR) analysis

The primer sequences employed were as follows: KIT (Forward: AGCAAATCCATCCCCACACC, Reverse: GCACCCAGGGTTTTCCCAAA), CSF1R(Forward: GGAGAGGAACGTGTGTCCAG, Reverse: GGATTCCCTGACCATGCCAA), GAPDH (Forward: ACAGCCTCAAGATCATCAGC, Reverse: GGTCATGAGTCCTTCCACGAT). To extract total RNA from cultured cells, Rnafast200 (Fastagen, Shanghai, China) was utilized, and subsequent cDNA synthesis was achieved using HiScript II Q RT SuperMix for quantitative PCR (qPCR) (Vazyme, Nanjing, China). Realtime (RT)-qPCR was carried out using ChamQ Universal SYBR qPCR MasterMix (Vazyme) in accordance with the manufacture’s instructions. The RT-qPCR procedure encompassed the following steps: initial denaturation at 95℃ for 30 s (one cycle); denaturation at 95℃ for 10 s, followed by 40 cycles; and a melting curve analysis at 95℃ for 15 s, 60℃ for 60 s and 95℃ for 15 s (one cycle). The quantification of gene expression levels was normalized to GAPDH and computed utilizing the log 2–44Ct method.

### 2.15 Chondrocytes isolation and culture

Chondrocytes were obtained using methods described in our previous study[30]. In brief, the articular cartilage of the OA patients was harvested and digested with 0.2% collagenase type II (Sigma, USA) at 37°C for 4 hours. Cells were cultured in DMEM/F-12 (Gibco, USA) supplemented with 10% FBS (Gibco, USA) at 37°C. Cells in passage 3 were used for co-culture. Furthermore, we characterized chondrocyte morphology and proliferation under the microscope. The cells from each patient were not pooled at the time of the co-culture experiment.

### 2.16 Immunofluorescence Staining of cells

Chondrocytes are fixed with paraformaldehyde, and permeabilized with Triton X-100. After blocking, primary antibodies against KIT and CSF1R (abcam, UK) are applied. Secondary antibodies were detected using a fluorescent secondary antibody (Proteintech, China) or Rabbit streptavidin-biotin detection system kit (ZSGB-Bio, China) according to the manufacturer’s protocol, followed by nuclear staining with DAPI. The cells were counted in five random fields per well. The percentage of positive cells was calculated using Image-Pro Plus version 6.0 for Windows (Media Cybernetics, USA).

### 2.17 Western blotting

Protein extracts were subjected to sodium dodecyl sulfate-polyacrylamide gel electrophoresis (SDS-PAGE), and the proteins were transferred to polyvinylidene fluoride (PVDF) membranes and blocked in blocking buffer (5% skimmed milk) for 1 hour. The membranes were incubated overnight at 4°C with primary antibodies against KIT, GSF1R and glyceraldehyde-3-phosphate dehydrogenase (GAPDH). The following day, the membranes were incubated with fluorophore-conjugated secondary antibody or HRP-conjugated secondary antibody at room temperature for 1 hour and developed in electrochemiluminescence (ECL) Western blot detection reagents (Biosharp, China). The band was analyzed using UVP Chem studio PLUS 815 (Analytik Jena, Germany).

### 2.18 Statistical analysis

All analyses were applied utilizing the R software (v 4.2.2). The Wilcoxon test was applied to determine the differences between the OA and control groups (p < 0.05).

## 3. Results

### 3.1 Identification and function analysis of candidate genes

In the training dataset, compared with the control group, 5,277 DEGs were identified in the OA group, including 2,588 up- and 2,689 down-regulated genes. Top 10 up-regulated DEGs include ASPM, KCNJ6, DIAPH3, BUB1B, COL1A1, NELK, CDH2, CDH10, CEP55, and POSTN; while top 10 downregulated DEGs include ADM, DUSP2, FOSB, PND1, ASTL, OPRK1, MYOC, SERPINA2, PLIN5, and CYP1A1 (**Figure 1A**-**B**). The intersection analysis among 5,277 DEGs, 238 RSPRGs, and 202 HSPRGs yielded 11 candidate genes (**Figure 1C)**.

**Figure 1.**
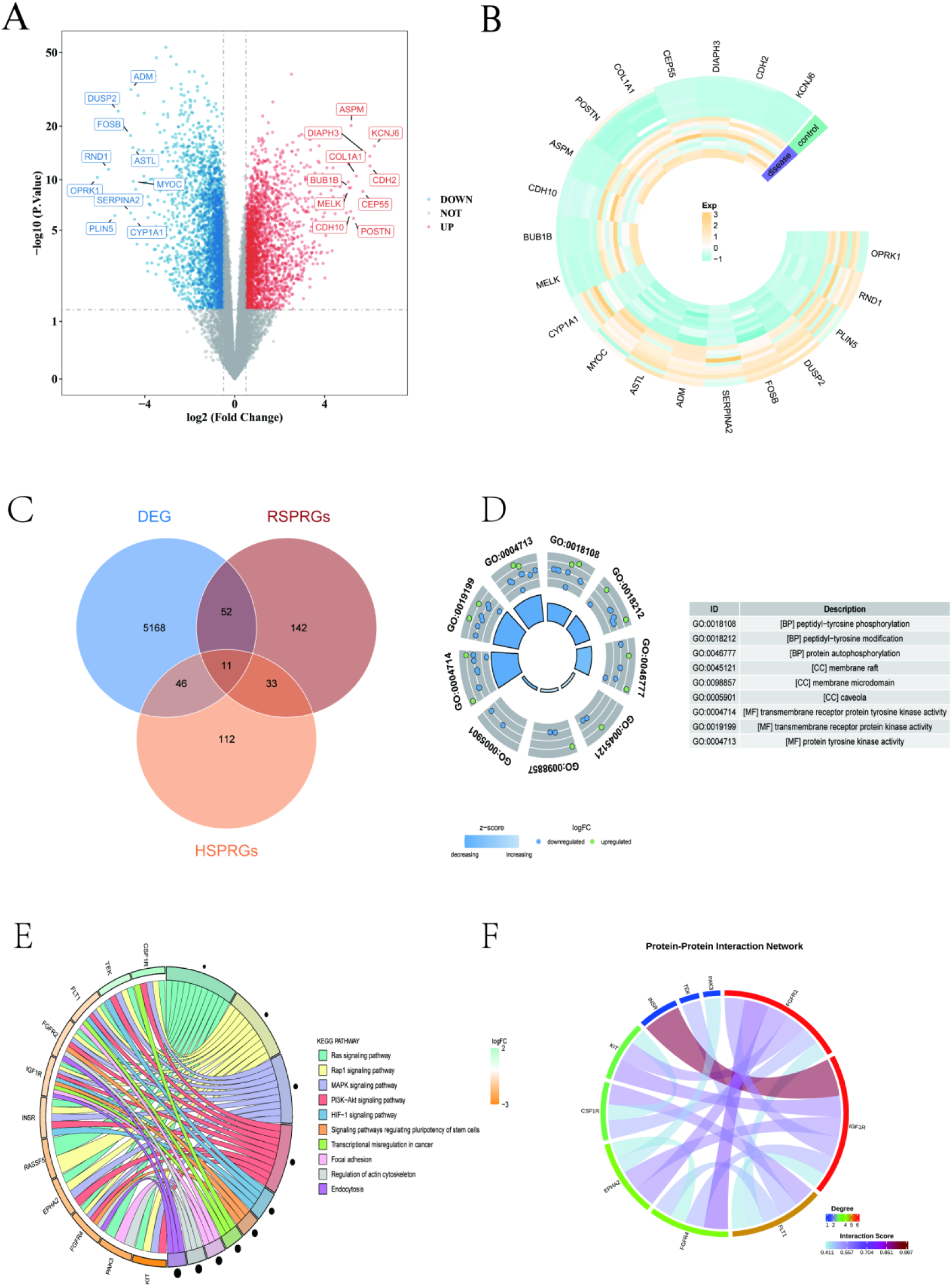
Identification and functional analysis of candidate genes in osteoarthritis. (A) Volcano plot of differentially expressed genes (DEGs) between OA and control groups. (B) Heatmap of the top 10 up- and down-regulated DEGs. (C) Venn diagram showing the overlap among DEGs, RSPRGs, and HSPRGs. (D) Gene Ontology (GO) enrichment analysis of candidate genes. (E) Kyoto Encyclopedia of Genes and Genomes (KEGG) pathway enrichment analysis of candidate genes. (F) Protein-protein interaction (PPI) network of candidate genes.

The GO enrichment analysis demonstrated that 11 candidate genes were significantly enriched in 416 pathways, including 379 BPs, 8 CCs, and 29 MFs. Top 3 BPs (coranked by P.value) included peptidyl-tyrosine phosphorylation, peptidyl-tyrosine modification, and protein autophosphorylation; for CCs, top 3 enrichment pathways included membrane raft, membrane microdomain, and caveola; in MFs, top 3 enrichment pathways included transmembrane receptor protein tyrosine kinase activity, transmembrane receptor protein kinase activity, and protein tyrosine kinase activity (P.adj < 0.05) (**Figure 1D, Supplementary Table 3**). KEGG enrichment analysis identified significant enrichment in 27 pathways, the 11 candidate genes were revealed that the top 10 most significantly enriched pathways (coranked by P.value) were Ras signaling pathway, Rap1 signaling pathway, MAPK signaling pathway, PI3K-Akt signaling pathway, HIF-1 signaling pathway, signaling pathways regulating pluripotency of stem cells, transcriptional misregulation in cancer, Focal adhesion, regulation of actin cytoskeleton, and endocytosis (P. adj < 0.05) (**Figure 1E, Supplementary Table 4**). The PPI network comprised 10 protein nodes with 17 interaction relationships among them, with the exception of one outlier. Among these, FGFR2, IGF1R, and FLT1 showed the most frequent interactions (**Figure 1F**).

### 3.2 Screening of candidate key genes and analysis of their correlation with YAP

The LASSO regression analysis identified 8 feature genes (CSF1R, EPHA2, FGFR2, INSR, KIT, PAK3, RASSF5, and TEK) when lambda. min = 0.0772 (**Figure 2A-B**). In the SVM-RFE model, 9 feature genes (KIT, RASSF5, FGFR2, INSR, PAK3, EPHA2, FGFR4, IGF1R, and CSF1R) were identified when the model achieved the highest accuracy of 0.9 and the lowest error rate of 0.1 (**Figure 2C-D**). The XGBoost algorithm identified 8 feature genes (KIT, RASSF5, FGFR2, INSR, TEK, FGFR4, EPHA2, and CSF1R) based on their Gain values, ranked in descending order (**Figure 2E**). The intersection of feature genes obtained from three machine learning algorithms yielded 6 candidate key genes, including CSF1R, EPHA2, FGFR2, INSR, KIT, and RASSF5 (**Figure 2F**). The PPI network involving YAP and the candidate key genes consisted of six core nodes with six interaction edges, along with one outlier node. Among these, YAP exhibited potential interactions with EPHA2 and RASSF5 (**Figure 2G**). Correlation analysis between YAP and the candidate key genes revealed that YAP1 showed significant correlations with several candidate genes. Specifically, a strong positive correlation was observed between YAP1 and INSR (cor = 0.69, P < 0.001), and a moderate positive correlation was identified between YAP1 and KIT (cor = 0.48, P < 0.05), suggesting potential regulatory relationships between YAP1 and these candidate key genes (**Figure 2H**).

**Figure 2.**
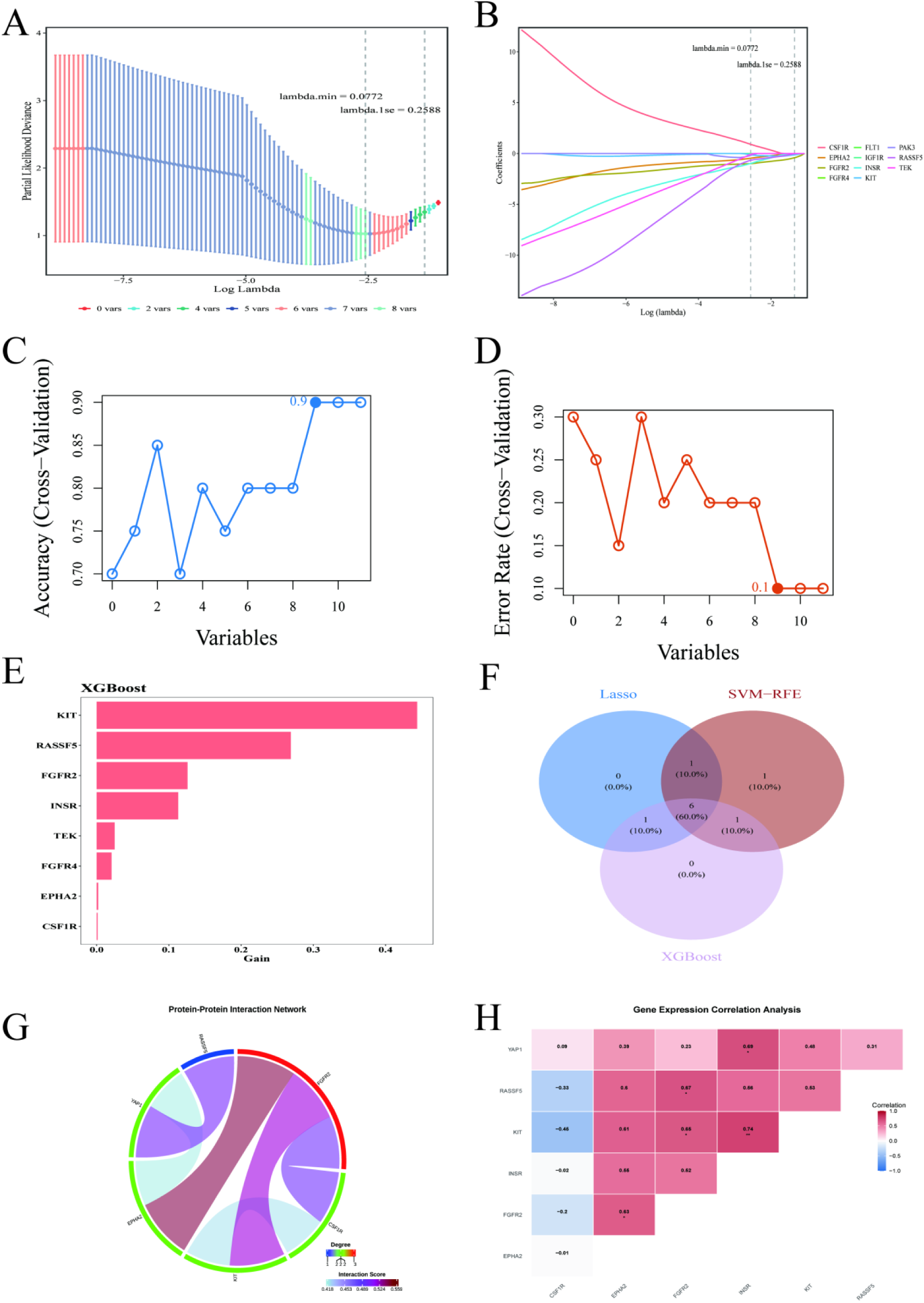
Screening of candidate key genes and their correlation with YAP. (A–B) Least absolute shrinkage and selection operator (LASSO) regression analysis with cross-validation curve and coefficient path. (C–D) Support vector machine-recursive feature elimination (SVM-RFE) algorithm performance and feature selection. (E) EXtreme Gradient Boosting (XGBoost) feature importance ranking. (F) Venn diagram of feature genes from three machine learning algorithms. (G) PPI network between candidate key genes and YAP. (H) Correlation heatmap between YAP and candidate key genes.

### 3.3 Identification of key genes and evaluation of their predictive efficacy

The expression patterns of CSF1R and KIT showed consistent trends between the training and validation datasets, with CSF1R significantly up-regulated and KIT significantly down-regulated in the OA group compared to the control group (P < 0.05) (**Figure 3A-B**). ROC analysis revealed that CSF1R (AUC = 0.770) and KIT (AUC = 0.940) exhibited AUC values exceeding 0.7 in the training dataset. These two genes were therefore established as key genes for OA (**Figure 3C-D**). According to the nomogram analysis, a higher total score was associated with an increased probability of OA. When the total score reached 173, the predicted probability of OA was 0.985 (**Figure 3E**). The calibration curve displayed a slope close to 1, and the HL test indicated good model fit (P = 0.133), demonstrating favorable predictive accuracy (**Figure 3F**). ROC analysis showed an AUC of 0.96, which exceeded the threshold of 0.7, demonstrating strong predictive performance of the model (**Figure 3G**). DCA further demonstrated a favorable net benefit across a wide range of threshold probabilities, indicating strong clinical utility of the nomogram model (**Figure 3H**).

**Figure 3.**
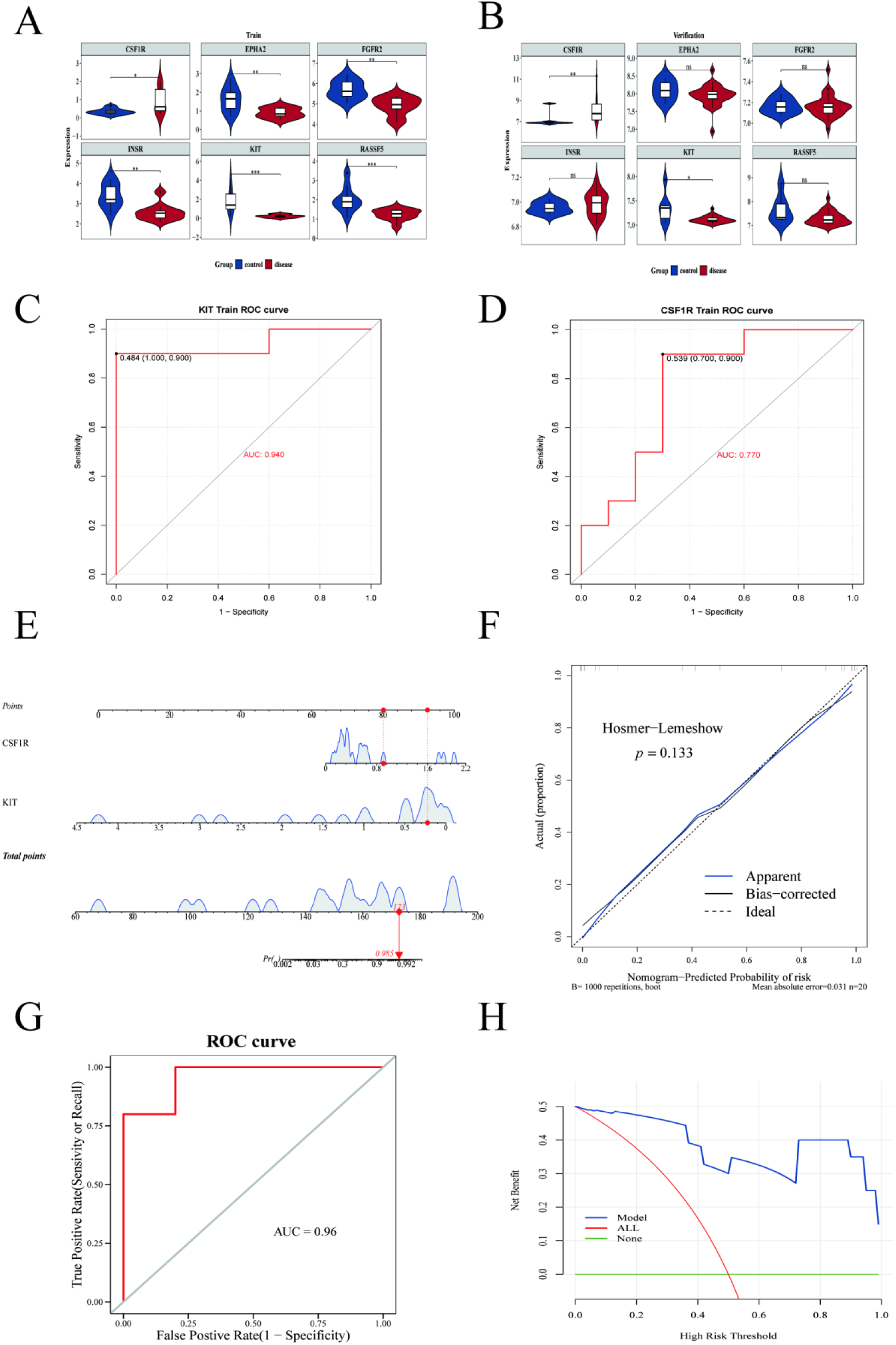
Identification of key genes and evaluation of their predictive efficacy. (A–B) Expression validation of key genes in training and validation datasets. (C–D) Receiver operating characteristic (ROC) curves of key genes in the training set. (E) Nomogram model for OA prediction. (F) Calibration curve of the nomogram. (G) ROC curve of the nomogram model. (H) Decision curve analysis (DCA) of the nomogram.

### 3.4 Interaction network, chromosomal mapping, and correlation of key genes

GeneMANIA analysis revealed that the top 5 genes exhibiting frequent interactions with the key genes CSF1R and KIT were PTPRU, CSF1, SOCS6, GRAP2, and SH2B3. The significantly enriched functional terms associated with these genes included receptor tyrosine kinase binding, peptidyl-tyrosine phosphorylation, cellular response to insulin stimulus, signaling receptor complex adaptor activity, and peptidyl-tyrosine modification (**Figure 4A**). Chromosomal localization analysis indicated that the KIT gene was located on chromosome 4, and the CSF1R gene was mapped to chromosome 5 (**Figure 4B**). Correlation analysis indicated a non-significant negative relationship between the two key genes (cor = −0.40, P = 0.083) (**Figure 4C**).

**Figure 4.**
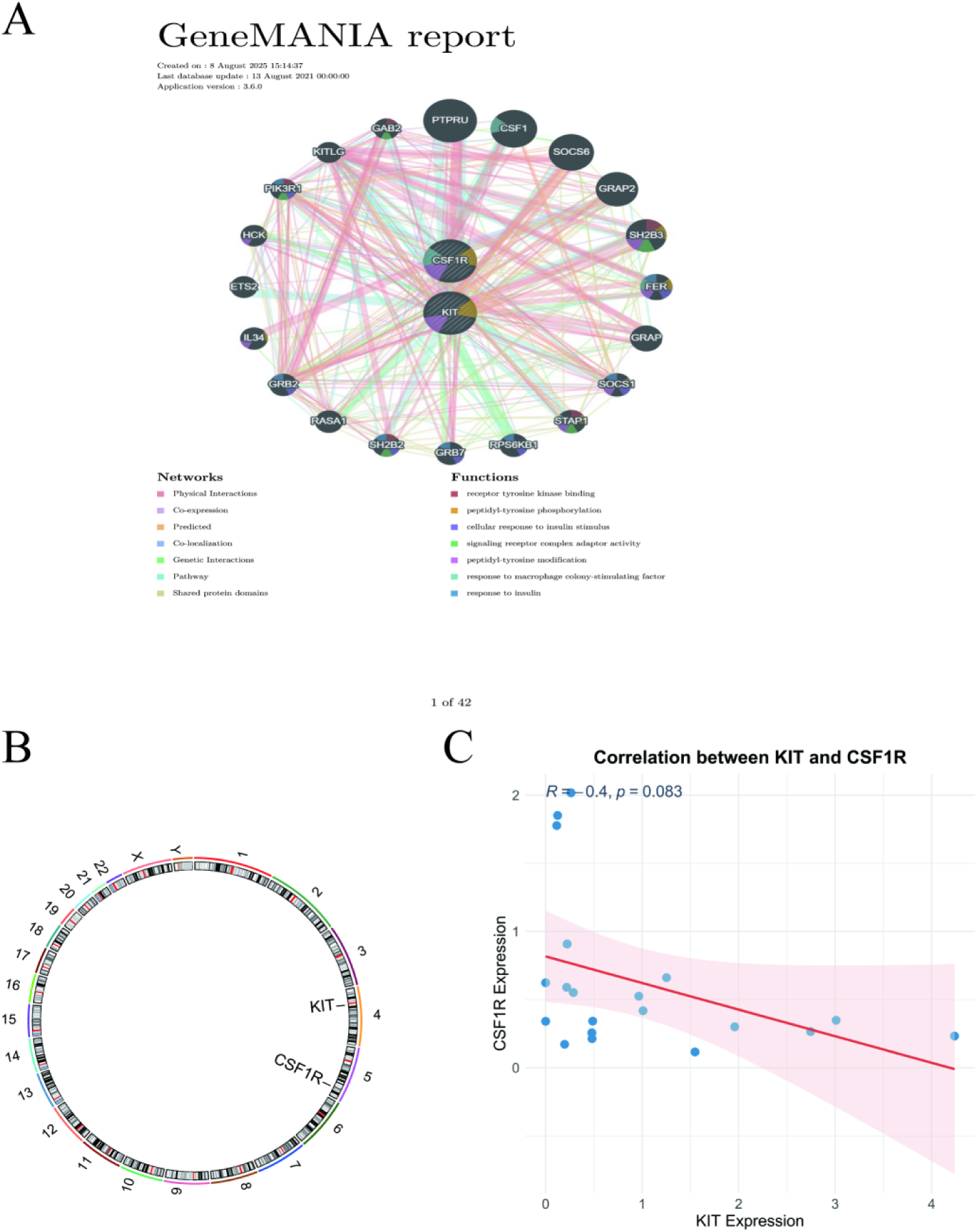
Interaction network, chromosomal mapping, and correlation of key genes. (A) GeneMANIA network showing functional associations of key genes. (B) Chromosomal localization of key genes. (C) Correlation scatter plot between key genes.

### 3.5 Enrichment pathways and subcellular localization analysis of key genes

GSEA results indicated that KIT was co-enriched in 11 pathways (|NES| > 1, P < 0.05, and P.adj < 0.25). When ranked by P-value, the 4 most significant pathways included ribosome, spliceosome, lysosome, and ECM-receptor interaction (**Figure 5A, Supplementary Table5**). Similarly, CSF1R was co-enriched in 9 pathways, with the top 4 being lysosome, focal adhesion, FcγR-mediated phagocytosis, and regulation of actin cytoskeleton (**Figure 5B, Supplementary Table6**). Subcellular localization analysis suggested that the protein products of KIT and CSF1R are likely localized to the nucleus, extracellular region, and plasma membrane (**Figure 5C-D**). These analyses systematically characterize molecular mechanisms associated with OA progression.

**Figure 5.**
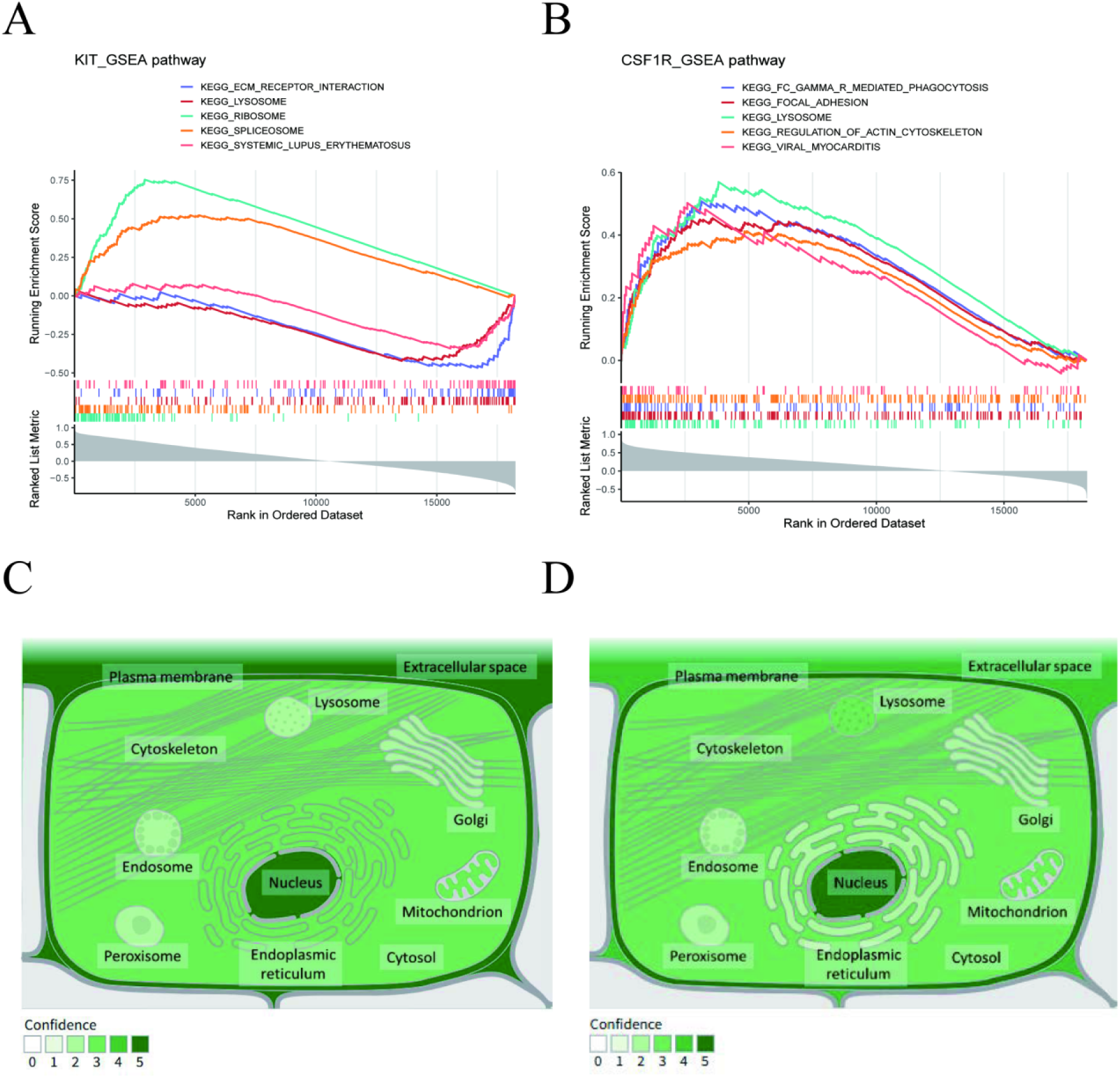
Enrichment pathways and subcellular localization of key genes. (A) Gene set enrichment analysis (GSEA) enrichment plot for KIT. (B) GSEA enrichment plot for CSF1R. (C-D) Subcellular localization predictions for KIT (C) and CSF1R (D).

### 3.6 Differential immune cells in OA

Immune cell infiltration analysis revealed significant differences in infiltration levels between the OA and control groups (P < 0.05). Among these, immature dendritic cells, central memory CD4⁺ T cells, and plasmacytoid dendritic cells exhibited relatively high infiltration levels in both groups (**Figure 6A**). Wilcoxon tests identified 14 differentially infiltrated immune cell types, including activated B cells, activated CD8⁺ T cells, CD56ᵇʳⁱᵍʰᵗ natural killer cells, effector memory CD8⁺ T cells, eosinophils, gamma delta T cells, immature dendritic cells, macrophages, mast cells, regulatory T cells, T follicular helper cells, type 17 T helper cells, and type 2 T helper cells (P < 0.05) (**Figure 6B**). Correlation analysis between key genes and differential immune cells demonstrated that KIT showed the most significant negative correlations with immature dendritic cells (cor = −0.823, P = 8.41 × 10⁻⁶), gamma delta T cells (cor = - 0.806, P = 1.76 × 10⁻⁵), and type 2 T helper cells (cor = −0.790, P = 3.45 × 10⁻⁵). CSF1R exhibited the strongest positive correlations with macrophages (cor = 0.814, P = 1.17 × 10⁻⁵), regulatory T cells (cor = 0.639, P = 0.0030), and T follicular helper cells (cor = 0.489, P = 0.0303) (**Figure 6C**). Correlation analysis among differentially immune cells indicated a highly positive correlation between T follicular helper cells and regulatory T cells (cor = 0.854, P < 0.001), and Type 2 T helper cells and immature dendritic cells also showed a strong correlation (cor = 0.833, P < 0.001). Significant positive correlations were observed between type 17 T helper cells and eosinophils (cor = 0.815, P < 0.001), regulatory T cells and macrophages (cor = 0.811, P < 0.001), and immature dendritic cells and gamma delta T cells (cor = 0.795, P < 0.001), suggesting potential cooperative roles in inflammatory and immune responses. Conversely, significant negative correlations were detected between immature dendritic cells and eosinophils (cor = −0.663, P = 0.0019), regulatory T cells and CD56ᵇʳⁱᵍʰᵗ natural killer cells (cor = - 0.669, P = 0.0017), regulatory T cells and eosinophils (cor = −0.686, P = 0.0012), gamma delta T cells and eosinophils (cor = −0.758, P < 0.001), and immature dendritic cells and activated B cells (cor = −0.774, P < 0.001), indicating potential antagonistic interactions within the immune microenvironment (**Figure 6D**). These findings reveal a complex and tightly regulated network among differential immune cells, suggesting that synergistic or antagonistic interactions may collectively drive OA progression and reshape the immune microenvironment.

**Figure 6.**
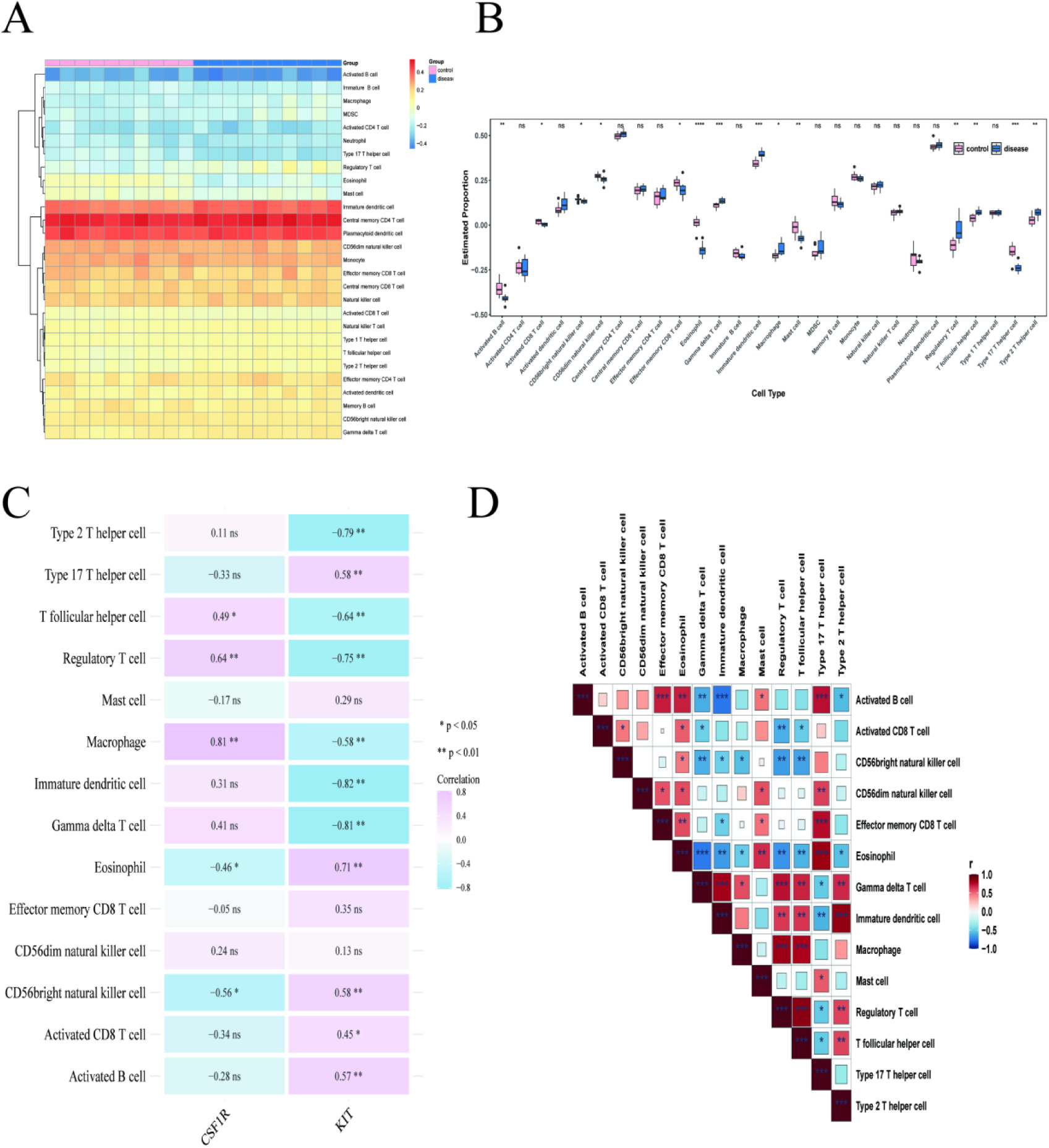
Differential immune cell infiltration in osteoarthritis. (A) Heatmap of immune cell infiltration levels. (B) Boxplot of differentially infiltrated immune cells. (C) Correlation heatmap between key genes and differential immune cells. (D) Correlation network among differential immune cells.

### 3.7 The relationship between key genes, regulatory factors, and potential drugs

The TF-miRNA-mRNA regulatory network comprised 45 nodes and 47 interaction edges. For KIT, 18 miRNAs and 11 TFs were predicted, while CSF1R was associated with 6 miRNAs and 12 TFs. Among these, 4 TFs (TFAP2A, STAT1, SMURF2, and WWP1) were commonly linked to both KIT and CSF1R (**Figure 7A**). In the drug- key gene interaction network, 115 nodes and 139 edges were identified. A total of 85 drugs were predicted for KIT and 54 for CSF1R, with 28 drugs shared between both genes, included Linifanib, AST-487, Cediranib, Mastinib, Pexidartinib, JNJ-7706621, Sorafenib, Entrectinib, SP 600125, Dovitinib, Sunitinib, BAY 61-3606, Dasatinib Anhydrous, Imatinib, GW843682X, Seralutinib, Ilorasertib, Pazopanib, Censertib, ENMD-2076, Pexidartinib Hydrochloride, Vatalanib, Tandutinib, RG-1530, CYC-116, and Quizartinib. The top 3 predicted drugs for KIT were Avapritinib, Bezclastinib, and Ripretinib, while those for CSF1R were IMC-CS4, Axatilimab, and Cabiralizumab (**Figure 7B**).

**Figure 7.**
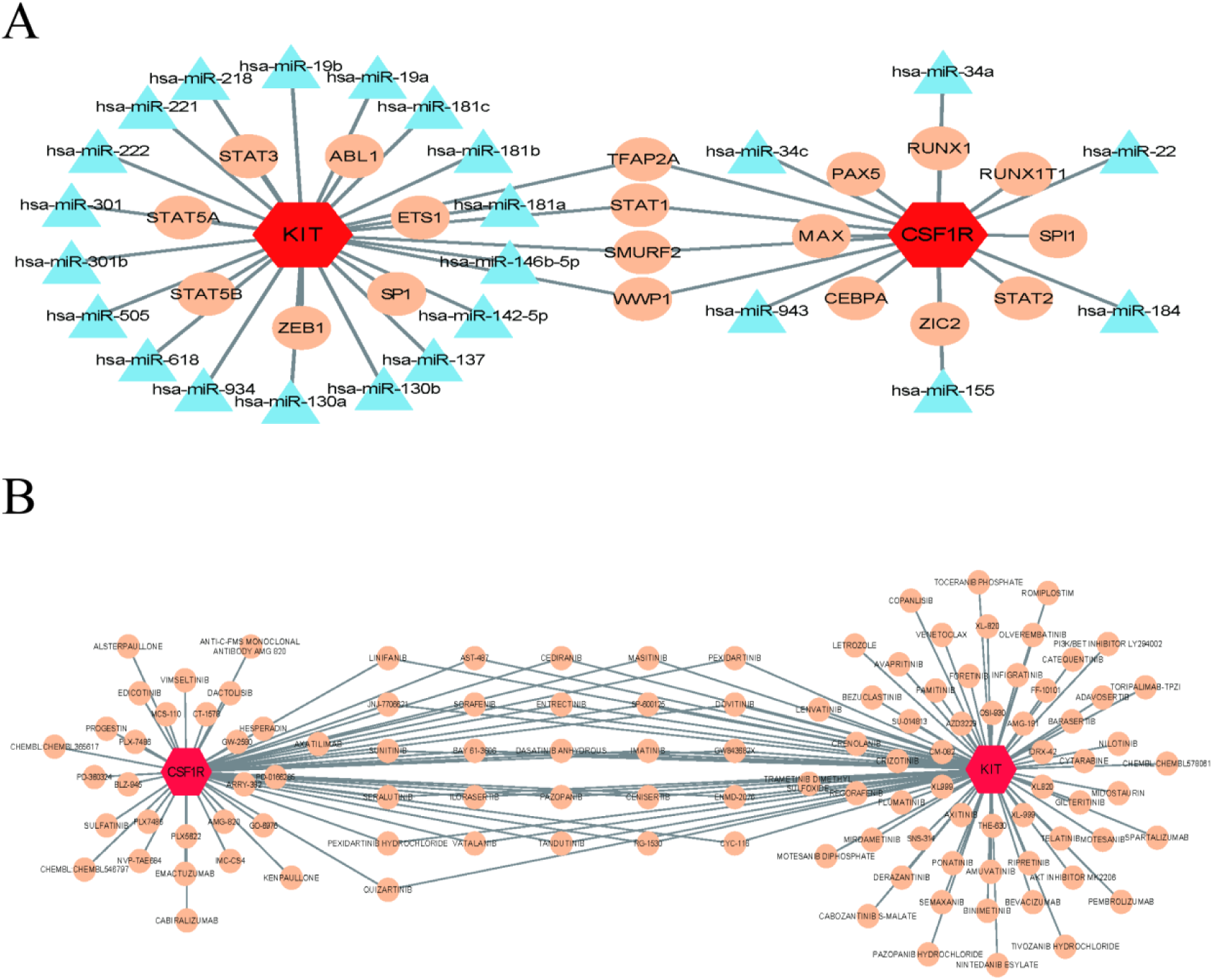
Regulatory network and drug prediction for key genes. (A) Transcription factor (TF)-microRNA (miRNA)-mRNA regulatory network. (B) Drug-key gene interaction network.

### 3.8 In Vitro Validation of key genes

To experimentally validate our bioinformatics findings, we performed real-time quantitative PCR and Western blot analysis on cartilage samples from osteoarthritis patients and healthy controls. Subsequently, we cultured chondrocytes from both groups and conducted immunofluorescence staining. Results revealed significantly upregulated CSF1R mRNA expression in OA tissue compared to controls (P < 0.0001), while KIT expression was significantly downregulated (P < 0.05, Figure 8A). Then we employed Western blot analysis to examine changes in protein levels within cartilage tissue from osteoarthritis patients and healthy controls. Compared to the control group, KIT protein levels were significantly reduced, while CSF1R protein expression levels were markedly elevated (Figure 8B). To further validate these findings, we examined the expression of KIT and CSF1R in chondrocytes. We observed the same trend: compared to the control group, OA chondrocytes exhibited significantly decreased KIT expression (P < 0.001, Figure 8C) and significantly increased CSF1R expression (P < 0.0001,Figure 8C). These findings align with our computational predictions, supporting the conclusion that CSF1R and KIT participate in osteoarthritis progression. This suggests CSF1R and KIT may serve as hallmark genes in the osteoarthritis process.

**Figure 8.**
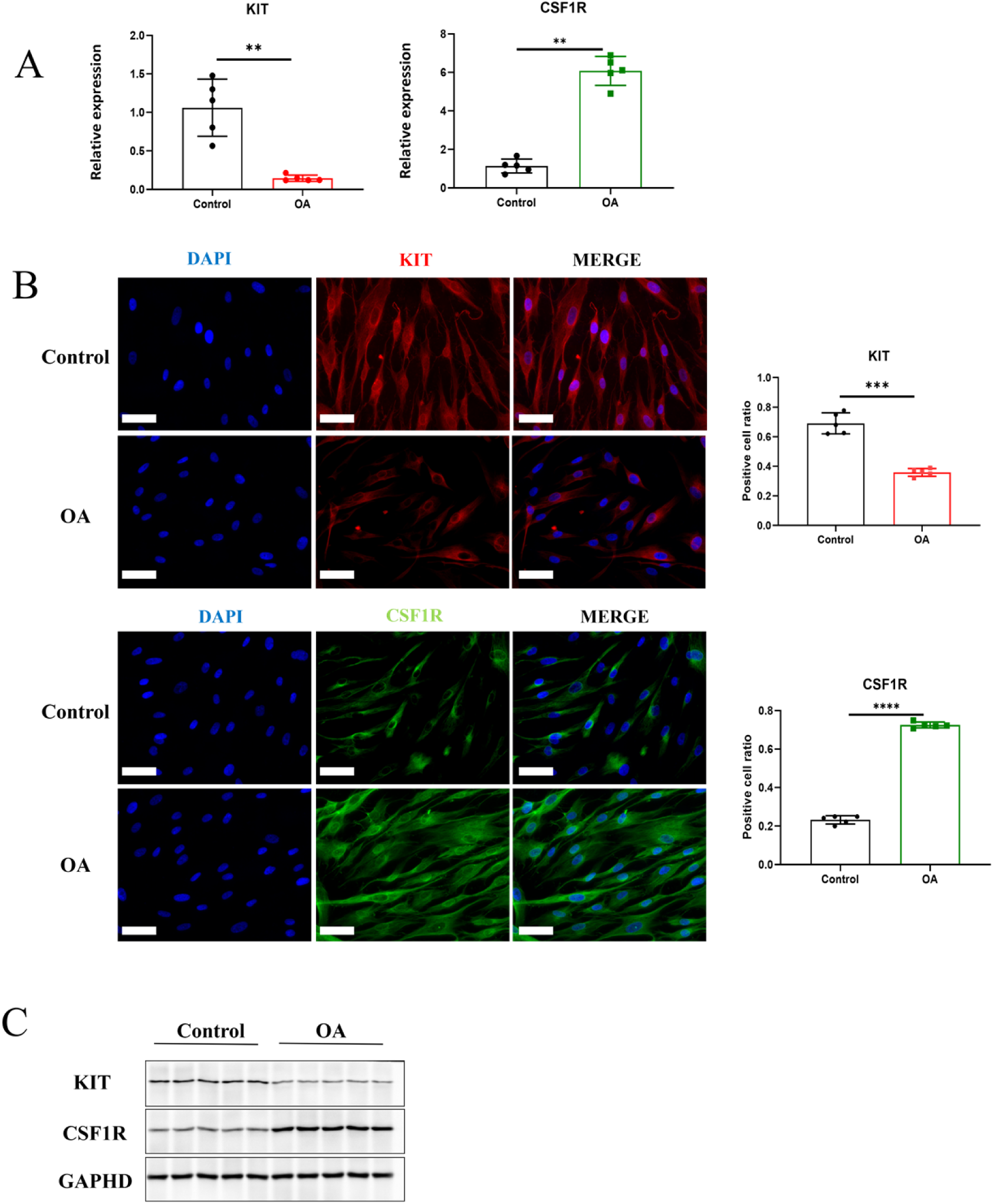
In vitro validation of key genes. (A)The mRNA expression of KIT, CSF1R in cartilage sample. (B) Immunohistochemistry staining of chondrocyte for KIT (red), CSF1R (green) and DAPI (blue). Bars=100μm. The cells were counted in five random fields per well. Bars = 100 μm. (C) The protein levels of KIT and CSF1R in cartilage sample. ***p < 0.001, ****p < 0.0001.

## Discussion

Osteoarthritis (OA) is a common degenerative joint disorder characterized by progressive articular cartilage erosion and subchondral bone remodeling [31]. Previous studies have demonstrated that the Ras signaling pathway contributes to OA pathogenesis by promoting synovial cell proliferation and regulating inflammation-mediated cartilage degradation, whereas the Hippo pathway maintains cartilage homeostasis by controlling chondrocyte proliferation and apoptosis [32,33]. Aberrant activation of the Ras signaling and the Hippo signaling may drive cartilage destruction and disease progression.

This study identified CSF1R and KIT as key genes in OA by integrating differential expression analysis with three machine learning algorithms. Expression validation and ROC analysis confirmed that CSF1R is significantly upregulated while KIT is significantly downregulated in OA, with both demonstrating good diagnostic performance. Functional analysis revealed KIT enrichment in metabolism-related pathways such as ribosomes and lysosomes, while CSF1R was associated with immune functions, including phagocytosis and cytoskeletal regulation. Immune infiltration analysis showed significant negative correlations between KIT and immature dendritic cells, and positive correlations between CSF1R and macrophages, suggesting their respective roles in OA progression through metabolic reprogramming and immune regulation. Regulatory network prediction identified four transcription factors, including STAT1, that jointly regulate both genes, and screened 28 potential targeted drugs. These findings systematically reveal the mechanism by which CSF1R/KIT acts as a pivotal node linking metabolism and immunity in OA.

KIT (stem cell factor receptor, SCFR; also known as CD117) encodes a type III transmembrane receptor tyrosine kinase that binds stem cell factor (SCF). In OA, KIT signaling primarily contributes to synovial inflammation. Elevated SCF expression in OA synovial tissue activates KIT, inducing ERK and STAT3 phosphorylation and promoting fibroblast-like synoviocytes to acquire a more destructive, pro-inflammatory phenotype [34]. This amplifies the release of inflammatory mediators and matrix-degrading enzymes, exacerbating cartilage loss. Elucidating the mechanisms of KIT activation in OA and developing targeted inhibitors may therefore represent novel therapeutic strategies for disease modification. GSEA further revealed the multidimensional functional network of KIT. Enrichment in ribosomal and spliceosomal pathways suggests that KIT-activated synovial cells exhibit a hypermetabolic state, supporting sustained high-level production of proinflammatory factors by enhancing mRNA processing and protein synthesis fidelity [35]. Enrichment in lysosomal pathways suggests KIT may regulate cellular stress responses by modulating the autophagy-lysosomal system, a mechanism particularly crucial under nutrient deprivation or inflammatory microenvironments [36]. More critically, enrichment of the ECM-receptor interaction pathway directly confirms KIT’s central role in cartilage matrix remodeling: activated KIT signaling not only promotes proinflammatory cytokine expression but may also amplify the positive feedback loop of matrix degradation by altering synovial cell adhesion to the ECM and signaling feedback [37].

CSF1R (colony-stimulating factor 1 receptor, also known as CD115 or c-FMS) is another member of the type III receptor tyrosine kinase family [38,39]. Binding of CSF1 or IL-34 to CSF1R regulates monocyte and macrophage proliferation, differentiation, and survival [40]. In OA, CSF1R is upregulated in synovial tissue and subchondral bone, where it mediates hypertrophic chondrocyte-induced inflammation and tissue remodeling[41,42]. CSF1R activation recruits and polarizes macrophages toward a pro-inflammatory phenotype, leading to overproduction of IL-1β, TNF-α, and matrix metalloproteinases (MMPs), which perpetuate cartilage degradation and synovial hyperplasia. Therefore, CSF1R serves as a molecular bridge linking subchondral bone remodeling and synovial inflammation in OA. GSEA analysis revealed the cellular biological basis of CSF1R function. Enrichment in the lysosomal pathway reflects a core feature of macrophage phagocytosis and degradation functions. CSF1R signaling enhances macrophage clearance of cartilage debris and apoptotic cells by regulating lysosomal biogenesis and function, but this process simultaneously releases proinflammatory factors to generate persistent inflammation[43]. Enrichment in the focal adhesion and actin cytoskeleton regulation pathways suggests CSF1R’s critical role in regulating macrophage adhesion, migration, and tissue infiltration [44,45] : Activated macrophages require dynamic reorganization of the actin cytoskeleton to traverse vascular barriers and localize to inflamed synovial and subchondral bone regions. The FcγR-mediated phagocytosis pathway further reinforces CSF1R’s central role in immune responses. This pathway, associated with antibody-immune complex clearance, suggests potential local immune complex deposition in OA synovium that activates macrophages [46].

Correlation analyses revealed a strong positive relationship between YAP (the Hippo pathway effector) and several Ras-associated genes, particularly KIT and INSR. This suggests that Hippo and Ras pathways engage in intricate crosstalk during OA progression, cooperatively influencing cell fate within the joint microenvironment [47]. YAP and its paralog TAZ act as mechanosensitive transcriptional co-activators that regulate chondrocyte proliferation, differentiation, and apoptosis in response to mechanical cues [48]. Our previous work also demonstrated that YAP regulates cartilage stem/progenitor cell (CSPC) function, thereby affecting OA progression. Notably, YAP correlates strongly with INSR, implicating potential cooperation with insulin/IGF signaling in modulating stem cell function [49,50]. Under abnormal mechanical stress, YAP may synergize with activated INSR signaling to impair stem cell homeostasis and accelerate cartilage degeneration. Furthermore, c-KIT can suppress MST1/LATS1 activity, leading to YAP dephosphorylation and nuclear translocation [51]. Thus, the YAP–Ras axis represents a potential integrative node and therapeutic target in OA [52].

Gene enrichment analysis (GESA) revealed that KIT is mainly enriched in ribosomal, spliceosomal, lysosomal, and ECM–receptor interaction pathways, suggesting roles in mRNA processing, protein synthesis fidelity, and extracellular matrix remodeling [53]. In contrast, CSF1R is enriched in lysosomes, focal adhesions, FcγR-mediated phagocytosis, and actin cytoskeleton regulation pathways [54], consistent with its regulatory role in macrophage adhesion, migration, and phagocytosis. The lysosomal pathway, as a common enrichment pathway for KIT and CSF1R, represents a critical intersection point for the functional convergence of these two genes. Although KIT primarily drives ECM degradation through synovial fibroblasts, while CSF1R mediates immune inflammation via macrophages, both genes rely on the lysosomal system to execute their respective effector functions. KIT may be involved in autophagy regulation and cellular stress adaptation, while CSF1R is directly linked to phagocytic degradation and antigen presentation. From a signaling integration perspective, direct molecular crosstalk exists between the Hippo-YAP pathway and mTORC1: LATS kinases inhibit mTORC1 activity by phosphorylating Raptor, while mTORC1 conversely regulates YAP degradation and nuclear localization by suppressing the autophagy-lysosomal system [55]. This bidirectional regulatory mechanism may occur at the lysosomal surface, where lysosomes serve not only as a key integration platform for mTORC1 signaling but also as a major convergence point for downstream effectors of the Ras-PI3K-AKT axis [56]. Thus, in the pathological context of osteoarthritis (OA), KIT and CSF1R—as common hub genes for Hippo and Ras pathways—promote catabolism and matrix degradation through activation of the Ras-mTORC1 axis on lysosomal membranes [57]. while the Hippo-YAP pathway influences lysosomal biogenesis and autophagic activity by regulating mTORC1 [58]. This functional complementarity suggests that simultaneous targeting of both molecules may yield synergistic therapeutic effects—inhibiting cartilage destruction while modulating the immune microenvironment—though experimental validation remains necessary.

Immune infiltration analysis showed that KIT expression negatively correlated with immature dendritic cells, γδ T cells, and Th2 cells, suggesting potential anti-inflammatory compensatory mechanisms despite its matrix-degrading effects.KIT expression negatively correlates with immature dendritic cells, γδ T cells, and Th2 cells. Immature DCs can transdifferentiate into functional osteoclasts in the presence of M-CSF and RANKL, a process significantly enhanced by synovial fluid from rheumatoid arthritis, potentially directly contributing to inflammatory bone lesions [59]. In rheumatoid arthritis and osteoarthritis studies, peripheral blood γδ T cells and Vδ2 T cells show significantly reduced proportions, with these cells migrating from peripheral blood to the synovium [60]. Th2 cells exert anti-inflammatory effects by secreting intrinsic anti-inflammatory cytokines such as IL-4, IL-5, IL-10, and IL-13, thereby delaying joint degeneration and cartilage damage [61]. This indicates that KIT, when normally expressed, can inhibit DC differentiation into osteoclasts, maintain the balance of immune cells between peripheral and local sites, and support effective Th2 responses. This reveals KIT as a potential central regulatory molecule linking bone destruction and immune dysregulation, providing new theoretical support for stratified diagnosis and precision treatment of OA.

Conversely, CSF1R exhibited strong positive correlations with macrophages, Treg cells, and Tfh cells. The CSF1/CSF1R axis is a canonical pathway for macrophage recruitment and activation [62]. Macrophage accumulation drives chronic low-grade inflammation in OA, while Treg dysfunction and Th17/Treg imbalance further exacerbate joint destruction [60,61]. These findings indicate that CSF1R may modulate immune imbalance and pathological bone remodeling in OA.

Transcription factor prediction identified STAT1 and TFAP2A as potential common regulators of KIT and CSF1R. STAT1 is a key effector of interferon signaling that mediates pro-inflammatory gene expression and can directly upregulate CSF1R, forming a positive feedback loop [63]. It’s predicted regulation of KIT reveals a novel mechanistic link between interferon signaling, inflammation, and stem cell factor pathways [64]. In the chronic inflammatory state of OA, persistently activated STAT1 may simultaneously drive CSF1R-mediated immune inflammation and KIT-mediated matrix degradation, forming a synergistic amplification effect of “immune-matrix destruction.” This mechanism suggests that STAT1 may serve as a key molecular switch linking the two pathways, whose excessive activation integrates the pathological effects of KIT and CSF1R into a unified tissue-destructive process.

Moreover, miR-146a and miR-34a were predicted to target CSF1R and KIT, respectively. miR-146a functions as a feedback regulator of inflammation by suppressing TRAF6 and IRAK1 to dampen NF-κB activation [65,66], whereas miR-34a acts as a tumor suppressor that restrains KIT-mediated proliferative signaling[67]. Dysregulation of these miRNAs in OA may therefore disrupt the fine balance of inflammatory and proliferative signaling, offering potential targets for miRNA-based interventions.

Drug–gene interaction analysis identified 28 compounds capable of co-targeting KIT and CSF1R, including masitinib, pexidartinib, sunitinib, and imatinib—all clinically approved or under investigation tyrosine kinase inhibitors (TKIs)[68]. Imatinib and sunitinib, for example, inhibit multiple RTKs, including KIT and PDGFR [69,70]. These agents theoretically could modulate both cartilage metabolism and macrophage-driven inflammation simultaneously [71]. Notably, pexidartinib—an FDA-approved CSF1R inhibitor—has shown efficacy in treating tenosynovial giant cell tumor (TGCT), a condition characterized by CSF1/CSF1R-mediated macrophage accumulation [72]. Given the mechanistic parallels, selective targeting of CSF1R may represent a feasible approach for OA subtypes with pronounced synovitis. Given the mechanistic similarities between TGCT and OA synovitis—both characterized by macrophage infiltration, chronic inflammation, and tissue destruction [73]—selective targeting of CSF1R may represent a viable therapeutic strategy for OA subtypes exhibiting a pronounced synovitis phenotype.

This study is based on integrative bioinformatic analyses, and functional validation remains necessary to determine causality. While the expression of KIT and CSF1R was independently verified, whether these genes act as drivers or consequences of OA pathology requires further investigation. Functional assays using gene knockdown or overexpression in chondrocytes, synoviocytes, and macrophages will be critical to confirm their roles. Furthermore, systemic administration of TKIs may cause off-target effects; thus, localized intra-articular delivery or nanoparticle formulations should be explored to enhance efficacy and reduce toxicity.

## Conclusion

Our findings identify KIT and CSF1R as key nodes linking the Ras and Hippo pathways, bridging inflammatory and mechanical signaling in OA. These results provide theoretical foundations and novel insights for developing targeted diagnostic and therapeutic strategies toward precision management of OA.

## Author Contributions

LZ: Acquisition and analysis of data. YL: Conceptual design and wrote the manuscript. DL: Data analysis. BS: Supervision of the entire study. All authors who contributed to the article and approved the submitted version.

## Data Availability Statement

The data that support the findings of this study are openly available in GSE114007 and GSE57218 at https://www.ncbi.nlm.nih.gov/geo/.

## Acknowledgments

Non

## Supplementary Table Legends

Supplementary Table 1. List of Ras signaling pathway-related genes (RSPRGs).

Supplementary Table 2. List of Hippo signaling pathway-related genes (HSPRGs).

Supplementary Table 3. Complete GO enrichment results for candidate genes.

Supplementary Table 4. Complete KEGG enrichment results for candidate genes.

Supplementary Table 5. GSEA enrichment results for KIT.

Supplementary Table 6. GSEA enrichment results for CSF1R.

